# SeqLengthPlot: An easy-to-use Python-based Tool for Visualizing and Retrieving Sequence Lengths from fasta files with a Tunable Splitting Point

**DOI:** 10.1101/2024.06.07.597948

**Authors:** Dany Domínguez-Pérez, Guillermin Agüero-Chapin, Serena Leone, Maria Vittoria Modica

## Abstract

**Motivation:** Accurate sequence length profiling is essential in bioinformatics, particularly in genomics and proteomics. Existing tools like SeqKit and the Trinity toolkit, among others provide basic sequence statistics but often fall short in offering comprehensive analytics and plotting options. For instance, SeqKit is a very complete and fast tool for sequence analyses, that delivers useful metrics (e.g., number of sequences, average, minimum, maximum length), and can returns the range of sequence shorter or longer (one side, not both at once) on a given lengths. Similarly, Trinity’s utility pearl-based scripts provide detailed contig length distributions (e.g., N50, median, and average lengths) but do not encompass the total number of sequences nor offer graphical representations of data.

**Results:** Given that key sequence analysis tasks are distributed among separate tools, we introduce SeqLengthPlot: an easy-to-use Python-based script that fills existing gaps in bioinformatics tools on sequence length profiling, crucial. SeqLengthPlot generates comprehensive statistical summaries, filtering and automatic sequences retriving from the input FASTA (nucleotide and proteins) file into two distinct files based on a tunable, user-defined sequence length, as well as the plots or dynamic visualizations of the corresponding sequences.

**Availability and implementation:** The detailed SeqLengthPlot pipeline is available on GitHub at https://github.com/danydguezperez/SeqLengthPlot, released under the GPL-3.0 license. Additional datasets used as sources or compiled as use cases are publicy available through the Mendeley Data repository: **DATASET_Ss_SE.1**: http://dx.doi.org/10.17632/pmxwfjyyvy.1, **DATASET_Ss_SE.2**: http://dx.doi.org/10.17632/3rtbr7c9s8.1, **DATASET_Ss_SE.3**: http://dx.doi.org/10.17632/wn5kbk5ryy.1, **DATASET_Ss_SE.4**: http://dx.doi.org/10.17632/sh79mdcm2c.1 and **DATASET_Ss_SE.5**: http://dx.doi.org/10.17632/zmvvff35dx.1.

## Introduction

Some tools are available for profiling sequence statistics on FASTA datasets and manipulation, such as those provided by the SeqKit tools (Shen *et al*., 2024, 2016), BigSeqKit (Piñeiro and Pichel, 2023), Biopython (Cock *et al*., 2009), BioPerl (Stajich *et al*., 2002), Pyfastx (Du *et al*., 2021), or others included in Seqfu (Telatin *et al*., 2021) or within Trinity’s utilities (Grabherr *et al*., 2011; Haas *et al*., 2013). These tools are commonly used to profile important statistics on de novo transcriptome assemblies. For instance, SeqKit and Seqfu comprises an extensive and complete set of useful tools, like the **seqkit stats** embedded in the SeqKit tool package delivering for one command the format (i.e., FASTA or FASTQ), type (DNA, RNA, protein), number of sequences, number of bases or residues, minimal, maximal and average sequence length for a given input FASTA file, while Trinity (i.e., **TrinityStats.pl)** and Seqfu provide detailed statistics on assembled-transcript length distribution, including contig length (N10-N50), median, and average length. Besides, **seqkit seq (-M or -m)** returns the range of sequence shorter or longer using a length threshold, but cannot deliver both splitted part of a given dataset at once, although it is possible to concatenate some functions in the same command line (i.e., **cat input_fasta** | **seqkit seq -m 100 -M 1000** | **seqkit stats**). However, these tools lack a visual output like plots, and thus, lack an all-in-one option to accomplish the above-mentioned tasks in a single run.

Commonly, 200 bp is the threshold for the *de novo* transcriptome assembly, while 100 amino acids is the standard when translating the corresponding Open Reading Frames (ORFs) with tools like TransDecoder (Haas and Papanicolaou, 2023). However, in both cases, the output might retain sequences below the set cutoff, requiring users to profile the translation accuracy (e.g., for NCBI submission) and sequence length distribution for proteomic studies. Likewise, retrieving sequences often requires additional steps using grep on Unix or a package like SeqKit. Considering that many of these tasks are provided in separated tools, herein, we introduce SeqLengthPlot.py version v1.0: a comprehensive and easy-to-use Python-based script that can handle many of these requirements in an unified tool. In this article, we showcase the implementation, requirements, outputs, and applications of SeqLengthPlot by testing it on original data derived from the single-end transcriptome of the false coral *Savalia savaglia* (Cnidaria).

### Tool Description

SeqLengthPlot.py version 1.0 is a straightforward Python-based script tailored for enhancing sequence length profiling. This script processes sequence data from a FASTA file to categorize and analyze transcript lengths. It generates histograms for transcript lengths above and below a specified threshold and provides statistical summaries of these distributions. The script also offers the flexibility to display or suppress plot pop-ups, making it suitable for both interactive analysis and automated pipelines.

### Main Components

- **Input Handling**: The script reads sequences from a specified nucleotide or protein FASTA file, which must be in the directory where the script is run or provided via an absolute path.
- **Output Directory Management**: By default, output files are saved in the same directory as the input FASTA file. Users have the option to uncomment a line in the script to instead save outputs directly to the home directory.
- **Length Thresholding**: Sequences are categorized based on a length threshold, set within the script to 200 base pairs. This threshold (**length_threshold**) is adjustable according to user needs.
- **Plot Generation**: Histograms of sequence lengths are generated for both above and below the threshold length, exporting linear and logarithmic scale plots. The popup display allow users to explore the plot interactively, and save it into .pdf, vectorized and different images formats (e.g., .png, .jpg, .tif, .eps, .pdf, .svg). To integrate into a pipeline, ensure the plot display lines are commented out.
- **Statistical Output**: The script calculates and saves statistics about the sequence lengths of the input and the resulting FASTA file, including total number of sequences, number of sequences above and below the threshold, and their minimum and maximum lengths.

### Applications

SeqLengthPlot is particularly useful for the straightforward assessment of:

- **Transcriptome Sequence Length Cutoff Accuracy:** Evaluate the accuracy of the standard cutoff length used by researcher when employing RNAseq assemblers such as Trinity through the abundance and distribution of transcripts shorter and longer than 200 base pairs (bp) (Fingerhut *et al*., 2018; Almeida *et al*., 2020). Since 200 bp is also the minimum required length for transcriptome submissions to databases like Transcriptome Shotgun Assembly (TSA, https://www.ncbi.nlm.nih.gov/genbank/tsa/), this assessment aids researchers in deciding whether to filter out shorter transcripts or retain them for potential overlapping re-assembly. By assessing SeqLengthPlot on original single-end transcriptome outputs from fragmented reads of Savalia savaglia (**DATASET_Ss_SE.1**), we demonstrate its value in a real-use case (**Fig. 1**).
- **Exploring ORF and Peptide Lengths for Biodiscovery:** Analyze the occurrence and distribution of ORFs generated by the Transdecoder (Haas and Papanicolaou, 2023) Supplementary **Fig. S1, DATASET_Ss_SE.2, DATASET_Ss_SE.4**), six-frame translation tool (Rice *et al*., 2000), Rapid Peptides Generator (Maillet, 2020), orfipy (Singh and Wurtele, 2021) or DeTox (Ringeval *et al*., 2024) output (Supplementary **Fig. S2, DATASET_Ss_SE.3, DATASET_Ss_SE.5**). This assessment enhances the accuracy, pre-functional classification and automatic retrieval of bioactive peptides such as antimicrobials peptides and animal toxins, mainly those that fall below or exceed common thresholds like 100 (Fingerhut *et al*., 2018; Almeida *et al*., 2020b; Agüero-Chapin *et al*., 2022), 60 (Barroso *et al*., 2024), 40 (Agüero-Chapin *et al*., 2023) or 30 amino acids (Hoepner *et al*., 2024), since certain algorithms are trained for specific peptide length ranges (Castillo-Mendieta *et al*., 2024; Rathore *et al*., 2023; Fingerhut *et al*., 2021; Santos-Júnior *et al*., 2020; Müller *et al*., 2017).

**Fig. 1:**
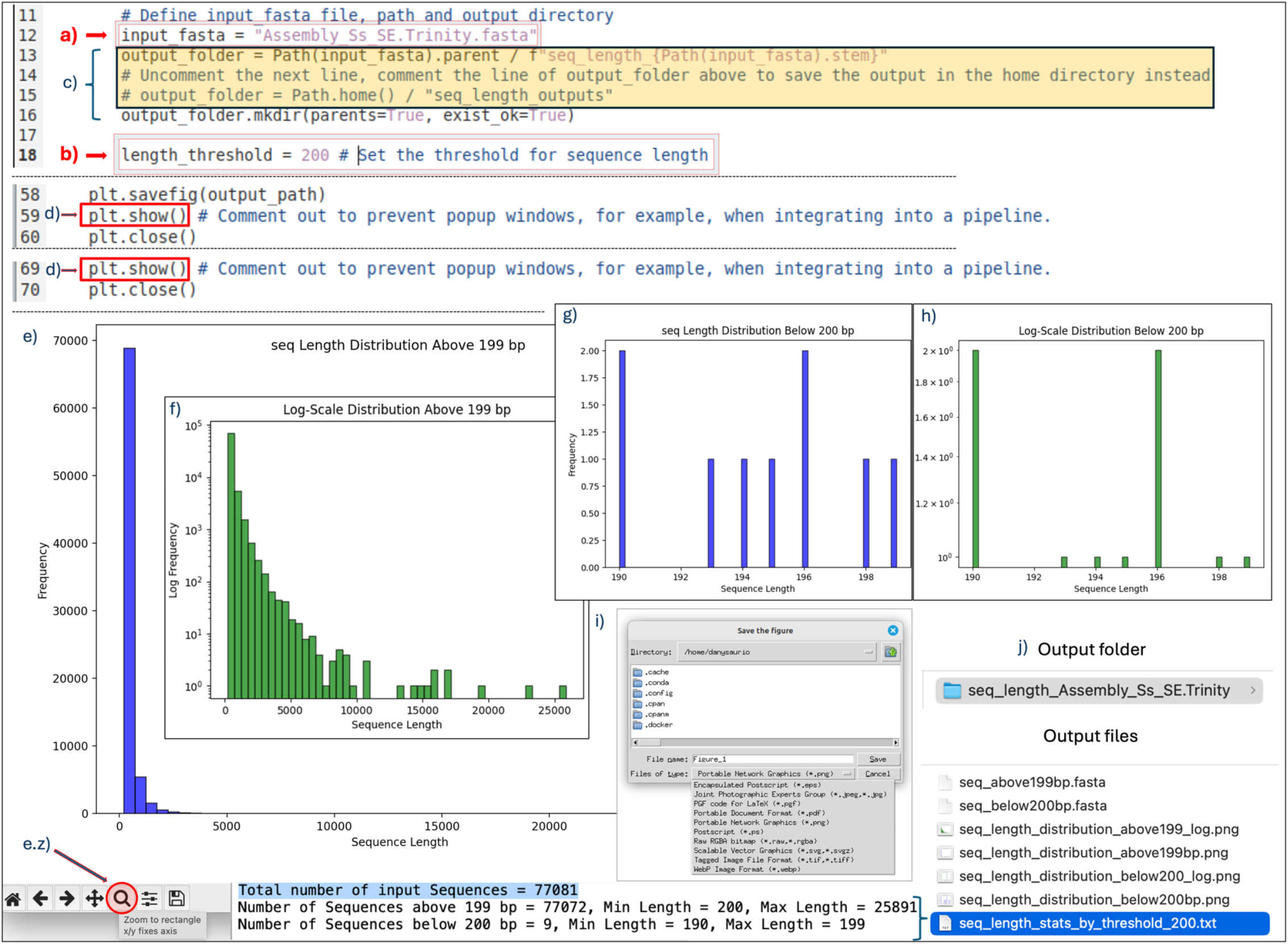
Illustrative diagram of the main components of SeqlenthPlot, depicting: a) path or name to the input_fasta file, b) sequence length cuttoff (default=200) to split the **input_fasta** file (i.e., **Assembly_Ss_SE.Trinity.fasta**), c) option to change the default output directory (parent directory of the input_fasta) to home, d) lines to comment out to prevent the plots from popping up (interactive plot set by default), e) linear-scale plot showing the sequences longer than 200 base-pairs (bp), e.z) zoom tool to explore the histogram (e.g., saturated areas of the plot), f) log-scale plot of the sequences longer than 200 bp enhancing the visibility of sequence distribution in the saturated length range of the histogram, g) linear and h) log-scale plots displaying sequences shorter than 200 bp (i.e., retained below the default length assembler “Trinity” cutoff), i) image format options to manually save the plots and j) output files and statistical summary.

### Compatibility, Installation and dependencies

This tool is compatible with both Unix and Windows operating systems. To run SeqLengthPlot.py, ensure you have an updated version of Python installed. Additionally, you will need the following libraries: matplotlib, Biopython, and Pathlib for plotting and sequence manipulation. Refer to our comprehensive guide on GitHub for detailed instructions on installing Python and dependencies across various configurations.

### Recommended implementation

Ensure your system is properly set up. Download the SeqLengthPlot.py script from https://github.com/danydguezperez/SeqLengthPlot and place it in a folder containing your input FASTA file. Open the script with a text editor and set the required parameters:

### Mandatory Parameters

a. **Define Path**: At **input_fasta =** “**Assembly_Ss_SE.Trinity.fasta**”, modify the path by replacing the default “**input_fasta**” file with “**your_path_or_your_input_fasta**” (**Fig. 1**a).
b. **Define the Sequence length:** At length_threshold = 200 (default length), set the threshold for your desired length cutoff (**Fig. 1**b).

### Optional Parameters

c) **Changing Output Path to home**: Users can comment out the defaults path **output_folder = Path(input_fasta).parent /** and uncomment (by removing the #) at the line **output_folder = Path.home() / “transcript_length_outputs”** to save the generated output-files in the home directory instead (**Fig. 1**c).
d) **Interactive Plots – Pipeline integration**: Users can comment out the **plt.show()** line to prevent plots from popping up. This is especially useful when integrating the script into automated data processing pipelines where no user interaction is desired (**Fig. 1**d). Plots will be saved anyway in the selected output directory.

### Running the script

Navigate to the folder containing **SeqLengthPlot.py** in the terminal or Command Prompt using the **cd** command. Then, execute the script in Unix systems by typing:

- **python3 SeqLengthPlot.py**

and in Windows.

- .**\SeqLengthPlot.py**

The script will generate files and plots automatically in a new folder named after your input FASTA file (**Fig. 1**j). If you encounter a “Warning: The system version of Tk is deprecated” message while plotting on MacOS, edit the script to switch the default backend from **matplotlib.use(‘TkAgg’)** to **‘MacOSX’** for interactive plots.

**Input File:**

- **Fasta Files**: nucleotides (e.g., “**Assembly_Ss_SE.Trinity.fasta**”) and proteins.

**Outputs files** (using sequence **length=200bp** by default):

- **Histogram Plots**: Four PNG files showing histograms of sequence lengths. Two are in linear scale (**seq_length_distribution_above199bp.png** and **seq_length_distribution_below200bp.png**), and two are in log scale (**seq_length_distribution_above199bp_log.png** and **seqs_length_distribution_below200bp_log.png**) (**Fig. 1**e-h).
- **Fasta Files**: Two fasta files (**seq_above199bp.fasta** and **seqs_below200bp.fasta**), splitted and retrieved from the orginal **input_fasta** file categorizing sequences based on the length threshold (**Fig.1**j).
- **Statistical Summary**: A text file (**seq_length_stats_by_threshold_200.txt**) containing detailed statistics of the sequence lengths on the **input_fasta**: Total number of input Sequences, Number of Sequences above 199 bp and below 200 bp, with the corresponding minimm and maximum lengths (**Fig.1**j).

## Conclusion

SeqLengthPlot.py merges multiple functionalities into a single, efficient platform, making it a useful and straightforward tool for enhancing sequence length distribution assessment on FASTA files, including plotting, filtering, and automatic sequence retrieval using a length threshold as a breaking point.

## Supporting information

SeqLengthPlot_Supplementary_Files

SeqLengthPlot_Supplementary_Data

## Acknowledgement

We thank Simonepietro Canese and Francesco Terlizzi from SZN for collecting the samples of *S. savaglia* used to generate the example data for this article. DDP acknowledge the support provided by the Centro Ricerche ed Infrastrutture Marine Avanzate in Calabria (CRIMAC) - Fondo FSC 2014-2020 - Piano Stralcio «Ricerca e Innovazione 2015-2017» – Programma Nazionale Infrastrutture di Ricerca (PNIR), CUP C64I20000320001.

## Notes

https://github.com/danydguezperez/SeqLengthPlot

http://dx.doi.org/10.17632/pmxwfjyyvy.1

http://dx.doi.org/10.17632/3rtbr7c9s8.1

http://dx.doi.org/10.17632/wn5kbk5ryy.1

http://dx.doi.org/10.17632/sh79mdcm2c.1

http://dx.doi.org/10.17632/zmvvff35dx.1

### Competing Interest Statement

The authors have declared no competing interest.

### Summary of Updates

In this version of the preprint, I have included the SeqLengthPlot Python script, along with the input and output files used for the analysis, to facilitate reproducibility of this previous version. Please note that a new version of SeqLengthPlot, introducing significant enhancements such as command-line functionality through flags, has been released alongside the corresponding peer-reviewed article (https://academic.oup.com/bioinformaticsadvances/article/5/1/vbae183/7905457). The GitHub repository has also been updated to reflect the latest version of the script. This update aims to avoid confusion for users cloning the repository and provides resources for testing the new version.

