## Supplementary material for "SeqLengthPlot: An easy-to-use Python-based Tool for Visualizing and Retrieving Sequence Lengths from fasta files with a Tunable Splitting Point": SeqLengthPlot_Supplementary_Files

- This file contains the Supplementary **Fig. S1** and **S2**, as well as additional graphical information to set up the SeqLengthPlot tool on real use cases

**Testing SeqLengthPlot.py on original single-end transcriptome outputs from the forward unpaired reads of *Savalia savaglia*'s RNAseq**

To test the SeqLengthPlot.py script on a real example, we selected the output files generated from the single-end transcriptome of the false coral *S. savaglia* using the remaining forward broken paired-end reads from the original RNAseq experiment (https://www.ncbi.nlm.nih.gov/bioproject/PRJNA1111802). We have previously used the resulting single-end transcriptome file **Assembly_Ss_SE.Trinity.fasta** as an illustrative example (**Fig. 1**). This file can be found as an input FASTA file of the SeqLengthPlot pipeline explanation on GitHub at https://github.com/danydguezperez/SeqLengthPlot, as well as can be downloaded from the Mendeley Data repository with other useful files generated as part of the transcriptomic workflow (**DATASET_Ss_SE.1**: http://dx.doi.org/10.17632/pmxwfjyyvy.1). Additionally, we provide two examples to demonstrate the application of SeqLengthPlot.py on real use cases and how to configure the script by adjusting the mandatory and optional parameters for ORFs obtained with TransDecoder v5.7.1 (Fig. S1, DATASET_Ss_SE.2: http://dx.doi.org/10.17632/3rtbr7c9s8.1, DATASET_Ss_SE.4: http://dx.doi.org/10.17632/sh79mdcm2c.1) and putative toxins identified by DeTox (Fig. S2, DATASET_Ss_SE.3: http://dx.doi.org/10.17632/wn5kbk5ryy.1, DATASET_Ss_SE.5: http://dx.doi.org/10.17632/zmvvff35dx.1) in the single-end transcriptome outputs from the forward unpaired reads of *S. savaglia* (DATASET_Ss_SE.1).

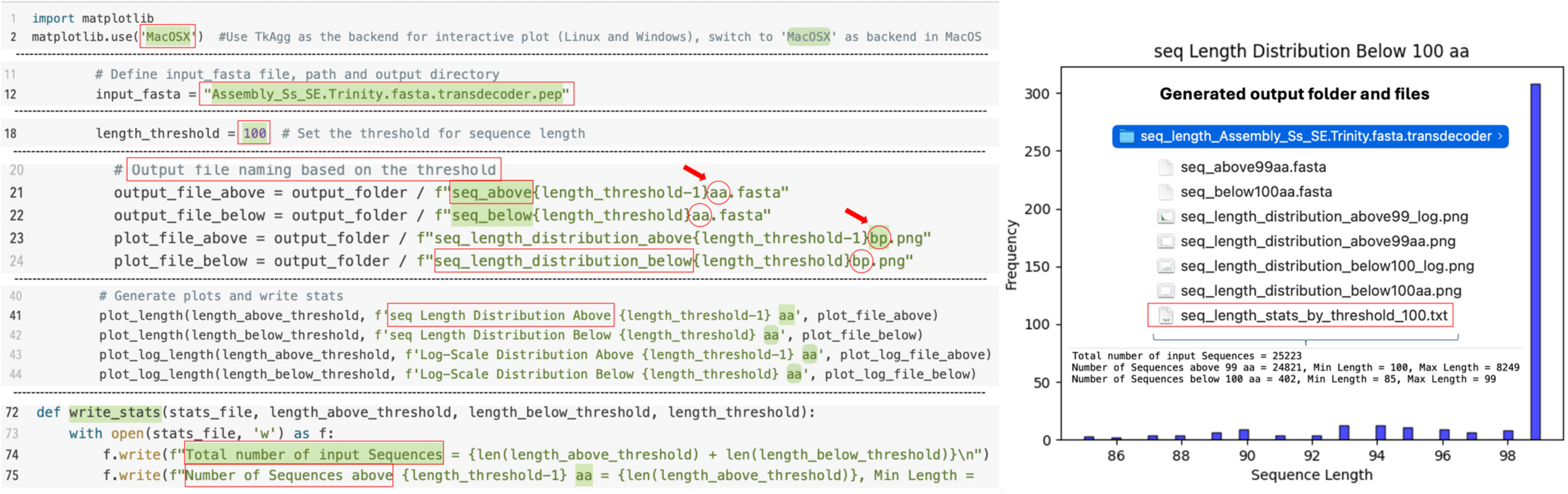


**Fig. S1**: Graphical illustration demonstrating the setup of SeqlengthPlot using a protein dataset obtained from TransDecoder and the resulting output files. The composite image highlights the flexibility to customize plot labels, legends, and statistical summaries by editing the SeqlengthPlot.py script. Specifically, in the left panel the **Line 2**: shows how to change the default backend used in Linux (**TkAgg**) to the recommended one for Mac users (**MacOSX**) to access the interactive plot options, **Line 12**: shows how to set up SeqLengthPlot.py by using a protein dataset as input FASTA file (**Assembly_Ss_SE.Trinity.fasta.transdecoder.pep**, **DATASET_Ss_SE.2**: http://dx.doi.org/10.17632/3rtbr7c9s8.1, **DATASET_Ss_SE.4**: http://dx.doi.org/10.17632/sh79mdcm2c.1), **Line 18**: defining sequence length cuttoff from (default=200) to 100, **Line 20 and Line 21**: comprises the sections to change legends of the plots and ouput files, highligthing the switch from base pair (bp) to amino acids (aa). Likewise, the section to write the statistics on **Line 72**, allow to customise the label of the statistics summary (**Line 74** and **Line 75**). The rigth panel shows the corresponding histogram plot in linear scale for the sequences below the selected threshold (100 aa), as well as the generated output folder, contained files, and written statistics according to the settings defined in the left panel.

```
11        # Define input_fasta file, path and output directory
12        input_fasta = "DeTox_output_Ss_SE_candidate_toxins.fasta"
13        #output_folder = Path(input_fasta).parent / f"seq_length_{Path(input_fasta).stem}"
14        # Uncomment the next line, comment the line of output_folder above to save the output in the home directory instead
15        output_folder = Path.home() / "seq_length_outputs"
16        output_folder.mkdir(parents=True, exist_ok=True)
17
18        length_threshold = 100  # Set the threshold for sequence length
-----------------------------------------------------------------------------------------------
52  def plot_length(data, title, output_path):
53      plt.figure()
54      plt.hist(data, bins=50, color='blue', edgecolor='black', alpha=0.7)
55      plt.xlabel('Sequence Length')
56      plt.ylabel('Frequency')
57      plt.title(title)
58      plt.savefig(output_path)
59      #plt.show() # Comment out to prevent popup windows, for example, when integrating into a pipeline.
60      plt.close()
-----------------------------------------------------------------------------------------------
69      #plt.show() # Comment out to prevent popup windows, for example, when integrating into a pipeline.
70      plt.close()
```

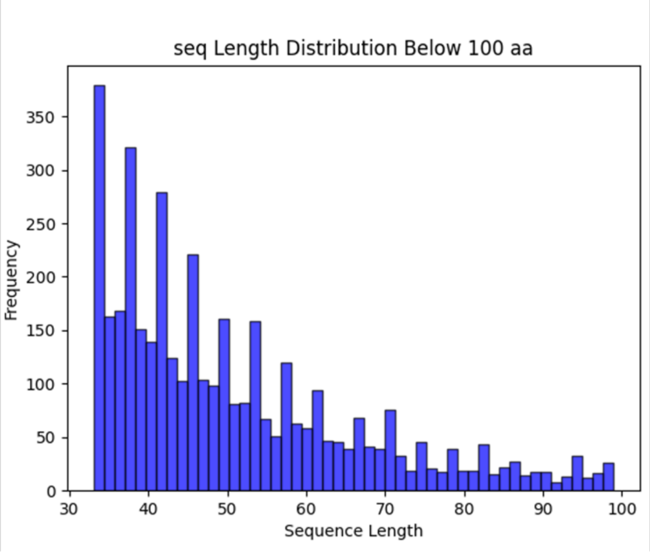


**Fig. S2**: Graphical illustration of setting up mandatory and optional parameters of SeqlengthPlot using a protein dataset obtained with DeTox and the generating plots. The left panel shows on **Line 12**: the input protein FASTA file obtained with the DeTox tool (**DeTox_output_Ss_SE_candidate_toxins.fasta**, **DATASET_Ss_SE.3**: http://dx.doi.org/10.17632/wn5kbk5ryy.1, **DATASET_Ss_SE.5**: http://dx.doi.org/10.17632/zmvvff35dx.1), provided in the path section (input_fasta). Lines 13-15 demonstrate how to change the default output directory from the parent directory of **input_fasta** to the home directory. Note: Unlike Linux, where changing this parameter will generate a folder named after your input FASTA file in your home directory, on a Mac, it will create a "seq_length_outputs" folder in the user path. The **Line 18**: defining sequence length cuttoff to 100 (default=200). By commenting out the **Lines 59** and **69** the user prevent the plots from popping up (interactive plot set by default). The rigth panel shows the corresponding histogram plot in linear scale for the sequences below the selected threshold (100 aa).

**Remarks on Key Features:**

- **Compatibility:** This tool is compatible with Unix and Windows Operating System (OS). It and has been tested on Linux (Linux Mint 21.3, Cinnamon 64-bits) and Windows 11 Pro (64-bits) using a High Performance Workstation (Lenovo ThinkStation P20, CPU: 2x 20-core Intel Gold 6148 CPU @ 2.40GHz, Linux Kernel: 5.15.0-105-generic, NVIDIA Quadro K620 2GB DDR3, 256 GB 2400 MHz DDR4, SSD NVMe KINGSTON SNV2S/2000G (2 TB - M.2 - 3500 MB/s), and on MacOS (Sonoma version 14.4.1) configuration (MacBook Pro 15-inch, 2019; 2,3 GHz 8-Core Intel Core i9; Intel UHD Graphics 630 1536 MB; 16 GB 2400 MHz DDR4, Mac HD 500 GB - APPLE SSD AP0512M) on Mac Studio with M2 Ultra chip, 128GB of RAM, 1TB SSD, and a 60-core GPU.

- **Comprehensive Metrics and Output**: Unlike seqkit stats, or TrinityStats.pl, SeqLengthPlot offers in a single tool the total number of sequences, minimum, maximum, of the input and splitted files, as well as the resulting FASTA files, containing the corresponding sequences length below and above the cuttoff.

- **Visual Analysis**: It generates intuitive plots for sequence length distributions, offering both linear and logarithmic views to accommodate a wide range of sequence lengths, enhancing data interpretation.

- **Flexible Threshold Settings**: The tool allows users to set custom length thresholds, crucial for tasks such as validating de novo transcriptome assemblies or analyzing protein-coding regions in ORFs, peptidomics and bioactive peptides biodiscovery which may require different length criteria.

- **Ease of Integration**: Designed for flexibility, it can be seamlessly run independently as a standalone script, or incorporated into larger bioinformatics workflows, supporting both interactive explorations and automated pipelines.

**Acknowledgement**

We thank Simonepietro Canese and Francesco Terlizzi from SZN for collecting the samples of *S. savaglia* used to generate the example data for this article. DDP acknowledge the support provided by the Centro Ricerche ed Infrastrutture Marine Avanzate in Calabria (CRIMAC) - Fondo FSC 2014-2020 - Piano Stralcio «Ricerca e Innovazione 2015-2017» – Programma Nazionale Infrastrutture di Ricerca (PNIR), CUP C64I20000320001.
