## Supplementary material for "SeqLengthPlot: An easy-to-use Python-based Tool for Visualizing and Retrieving Sequence Lengths from fasta files with a Tunable Splitting Point": SeqLengthPlot_Supplementary_Data: SeqLengthPlot.py

```
import matplotlib
matplotlib.use('TkAgg')  #Use TkAgg as the backend for interactive plot (Linux and Windows), switch to 'MacOSX' as backend in MacOS

import matplotlib.pyplot as plt
from Bio import SeqIO
from pathlib import Path
import sys

def main():
    try:
        # Define input_fasta file, path and output directory
        input_fasta = "Assembly_Ss_SE.Trinity.fasta"
        output_folder = Path(input_fasta).parent / f"seq_length_{Path(input_fasta).stem}"
        # Uncomment the next line, comment the line of output_folder above to save the output in the home directory instead
        # output_folder = Path.home() / "seq_length_outputs"
        output_folder.mkdir(parents=True, exist_ok=True)

        length_threshold = 200  # Set the threshold for sequence length

        # Output file naming based on the threshold
        output_file_above = output_folder / f"seq_above{length_threshold-1}bp.fasta"
        output_file_below = output_folder / f"seq_below{length_threshold}bp.fasta"
        plot_file_above = output_folder / f"seq_length_distribution_above{length_threshold-1}bp.png"
        plot_file_below = output_folder / f"seq_length_distribution_below{length_threshold}bp.png"
        plot_log_file_above = output_folder / f"seq_length_distribution_above{length_threshold-1}_log.png"
        plot_log_file_below = output_folder / f"seq_length_distribution_below{length_threshold}_log.png"
        stats_file = output_folder / f"seq_length_stats_by_threshold_{length_threshold}.txt"

        # Read and process sequences
        length_above_threshold, length_below_threshold = [], []
        with open(output_file_above, 'w') as file_above, open(output_file_below, 'w') as file_below:
            for record in SeqIO.parse(input_fasta, "fasta"):
                if len(record.seq) >= length_threshold:
                    SeqIO.write(record, file_above, "fasta")
                    length_above_threshold.append(len(record.seq))
                else:
                    SeqIO.write(record, file_below, "fasta")
                    length_below_threshold.append(len(record.seq))

        # Generate plots and write stats
        plot_length(length_above_threshold, f'seq Length Distribution Above {length_threshold-1} bp', plot_file_above)
        plot_length(length_below_threshold, f'seq Length Distribution Below {length_threshold} bp', plot_file_below)
        plot_log_length(length_above_threshold, f'Log-Scale Distribution Above {length_threshold-1} bp', plot_log_file_above)
        plot_log_length(length_below_threshold, f'Log-Scale Distribution Below {length_threshold} bp', plot_log_file_below)
        
        write_stats(stats_file, length_above_threshold, length_below_threshold, length_threshold)

    except Exception as e:
        print(f"An error occurred: {e}")
        sys.exit(1)

def plot_length(data, title, output_path):
    plt.figure()
    plt.hist(data, bins=50, color='blue', edgecolor='black', alpha=0.7)
    plt.xlabel('Sequence Length')
    plt.ylabel('Frequency')
    plt.title(title)
    plt.savefig(output_path)
    plt.show() # Comment out to prevent popup windows, for example, when integrating into a pipeline.
    plt.close()

def plot_log_length(data, title, output_path):
    plt.figure()
    plt.hist(data, bins=50, color='green', edgecolor='black', alpha=0.7, log=True)
    plt.xlabel('Sequence Length')
    plt.ylabel('Log Frequency')
    plt.title(title)
    plt.savefig(output_path)
    plt.show() # Comment out to prevent popup windows, for example, when integrating into a pipeline.
    plt.close()

def write_stats(stats_file, length_above_threshold, length_below_threshold, length_threshold):
    with open(stats_file, 'w') as f:
        f.write(f"Total number of input Sequences = {len(length_above_threshold) + len(length_below_threshold)}\n")
        f.write(f"Number of Sequences above {length_threshold-1} bp = {len(length_above_threshold)}, Min Length = {min(length_above_threshold, default='N/A')}, Max Length = {max(length_above_threshold, default='N/A')}\n")
        f.write(f"Number of Sequences below {length_threshold} bp = {len(length_below_threshold)}, Min Length = {min(length_below_threshold, default='N/A')}, Max Length = {max(length_below_threshold, default='N/A')}\n")

if __name__ == "__main__":
    main()
```
