## Supplementary figures and images for "SeqLengthPlot: An easy-to-use Python-based Tool for Visualizing and Retrieving Sequence Lengths from fasta files with a Tunable Splitting Point"

### seq_length_distribution_above199_log.png

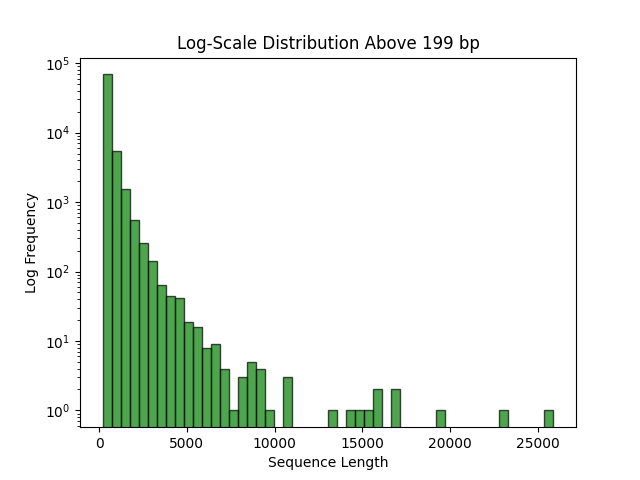

### seq_length_distribution_above199bp.png

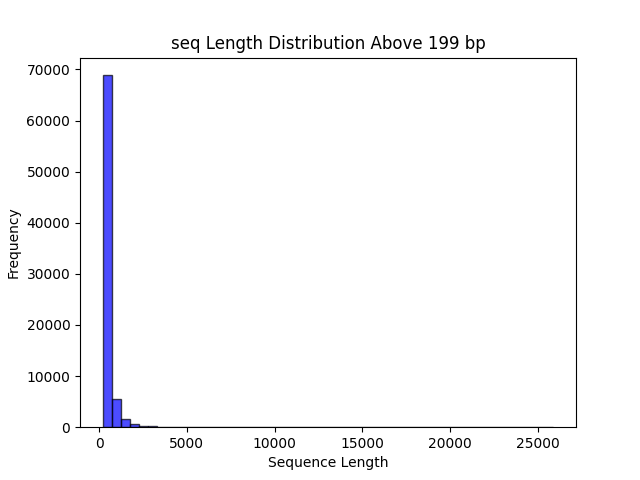

### seq_length_distribution_below200_log.png

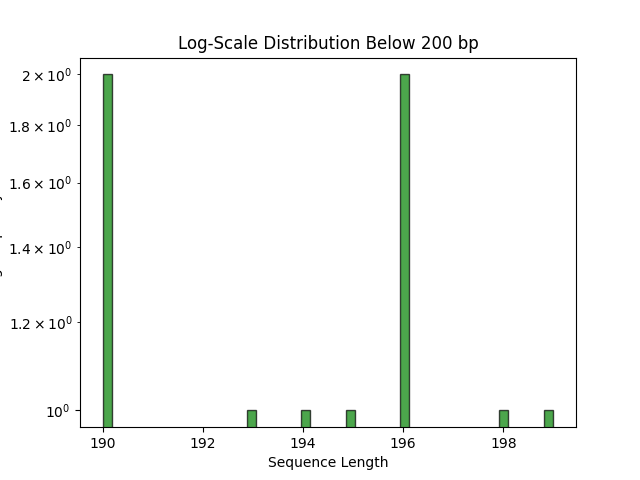

### seq_length_distribution_below200bp.png

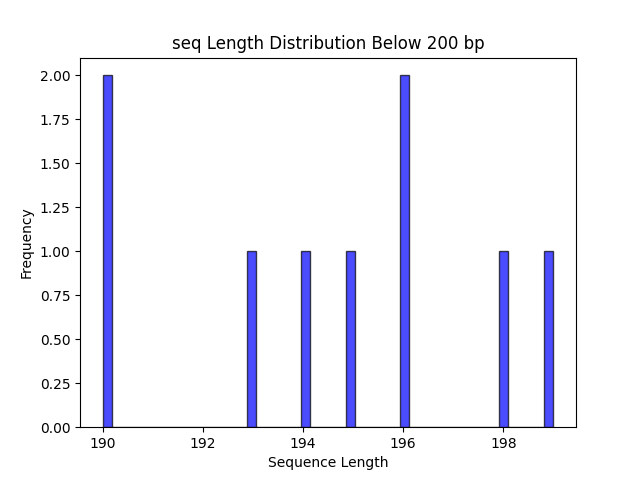
